# Reducing antimicrobial usage in small-scale chicken farms in Vietnam: A three-year intervention study

**DOI:** 10.1101/2020.09.13.295659

**Authors:** Doan Hoang Phu, Nguyen Van Cuong, Dinh Bao Truong, Bach Tuan Kiet, Vo Be Hien, Ho Thi Viet Thu, Lam Kim Yen, Nguyen Thi Tuyet Minh, Pawin Padungtod, Erry Setyawan, Guy Thwaites, Jonathan Rushton, Juan Carrique-Mas

## Abstract

Indiscriminate antimicrobial use (AMU) in animal production is a driver of antimicrobial resistance globally, with a need to define sustainable AMU-reducing interventions in small-scale farms typical of low- and middle-income countries. We conducted a before-and-after intervention study on a random sample of small-scale chicken farms in the Vietnamese Mekong Delta from 2016 to 2019. A baseline was established before providing farms (n=102) with veterinary advice on chicken health and husbandry, and antimicrobial replacement products. Thirty-five (34.2%) farms entered the intervention phase; the remainder no longer continued raising chickens. The intervention reduced AMU (−66%) (hazard ratio [HR]=0.34; p=0.002) (baseline 343.4 Animal Daily Doses per 1,000 chicken-days) and mortality (−40%) (HR=0.60; p=0.005) (weekly baseline 1.60 per 100). Chicken bodyweight increased by 100g (p=0.002) in intervention flocks. Our findings demonstrate that in the Vietnamese context, AMU can be substantially reduced in small-scale chicken farms without compromising flock health by providing veterinary advice.

## Introduction

In many low-and-middle income countries (LMICs) small-scale poultry farming plays a crucial role in supporting the livelihoods of rural communities (Wong et al., 2017). Compared with other species, poultry production has relatively low investment and production costs (Hilmi et al., 2011). Globally, poultry (mainly chicken) is the second most consumed type of meat (117 million tonnes in 2017), and by 2026 it is expected to surpass pork (OECD-FAO, 2017). Antimicrobial use (AMU) in animal production has been recognized as a driver of antimicrobial resistance (AMR) globally (Marshall & Levy, 2011; O’neill, 2015). In terms of frequency, chickens are the target of the highest AMU levels of all animal food species (Cuong et al., 2018). In addition, many antimicrobial active ingredients (AAI) regarded as critically important for human medicine by the World Health Organization (WHO, 2019) are often used in chicken production (Cuong et al., 2019).

In Vietnam, it has been estimated that three quarters (72%) of all AMU (3,842 tonnes in 2015) are aimed at animal production (Carrique-Mas et al., 2020). Studies in the Mekong Delta region of Vietnam have described very high amounts of antimicrobial to small-scale chicken flocks (Carrique-Mas et al., 2015; Trung et al., 2015; Cuong et al., 2019; Nhung et al., 2016). The high levels of disease and mortality in flocks in the area is a major driver of AMU in such systems (Carrique-Mas et al., 2019). In chicken farms, antimicrobials are used primarily for disease prevention (Carrique-Mas et al. 2015), since farmers regard them as a cheaper alternative to other disease control measures (Truong et al., 2019). Recent studies have shown that some of the most commonly used AAIs in small-scale chicken flocks in the area also belong to the WHO highest critical importance category such as polymyxins and fluoroquinolones (Cuong et al., 2019; Nhung et al., 2016). This situation is aggravated by a general lack of awareness about antimicrobials and the negative consequences of AMR among farmers (Pham-Duc et al., 2019). In addition, the ease of access to antimicrobials over the counter in veterinary drug shops (Phu et al., 2019) and their affordability (Dung et al., 2020) are factors that contribute to excessive AMU in Vietnam.

There is a pressing need to identify sustainable interventions that reduce AMU in food animal production systems. Such interventions will need to overcome the diversity of production systems and value chains they depend on and the patterns of AMU in these systems and policy contexts. A number of interventions have already taken place in developed countries based on improvements in biosecurity and husbandry practices aiming at reducing AMU in pigs (Raasch et al., 2020; Postma et al., 2017; Rojo-Gimeno et al., 2016) and broilers (Roskam et al., 2019).

However, no intervention studies targetting AMU in small-scale farming systems from LMICs have been published. We conducted a ‘before-and-after’ randomized intervention study on small-scale chicken farms in the Mekong Delta region of Vietnam. The intervention consisted on providing farmers with regular veterinary advice, alongside antimicrobial replacement products (Carrique-Mas & Rushton, 2017). The aim was to investigate the impact of this intervention on AMU, as well as on flock disease and productivity. Results and the lessons from this study can be adapted to comparable animal production systems in Vietnam and more generally, to other LMICs.

## Results

### Recruitment of study farms

The study took place between October 2016 and November 2019. A meeting with 199 randomly selected farmers from the farm census registered as owners of chickens was held in October 2016. Eighty-eight participating farmers indicating their willingness to restock within 6 months were enrolled. The remaining 14 farms were identified by commune animal health workers or through contact with farmers that had already been enrolled in the study. Therefore, a total of 102 farms were enrolled over the period October 2016 to October 2017. The baseline phase spanned October 2016 to April 2018. The intervention was delivered from May 2018 to November 2019.

The flow of participating farms was complicated by many (n=63) that stopped farming during the study for financial reasons unrelated to the study. The recruitment and allocation to arms is summarized in Figure 1. Their location is presented in Supplementary file 1.

**Figure 1.**
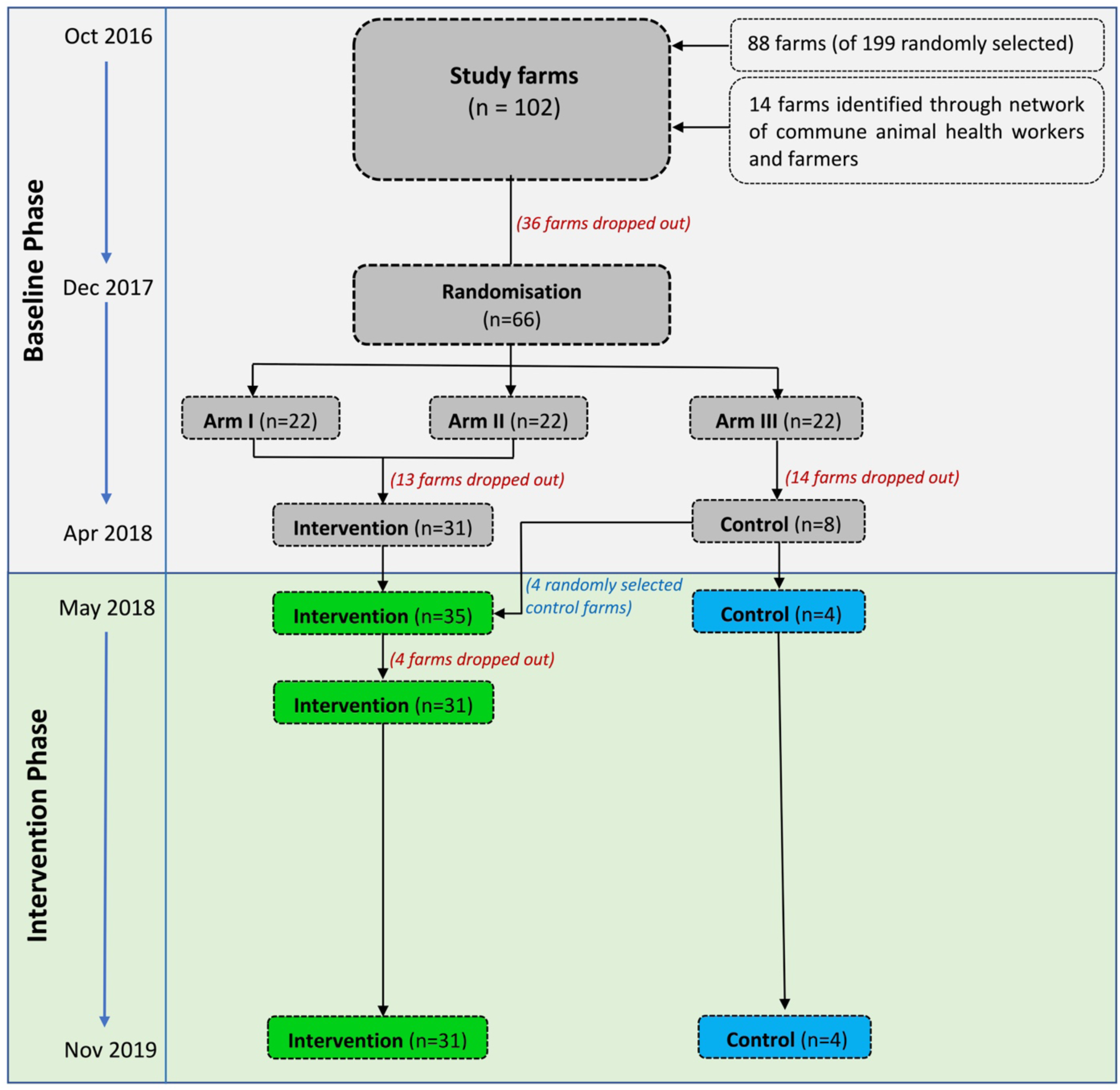
Recruitment and follow-up of study farms.

In December 2017 farms that remained in production at the time (n=66) were randomised to either intervention Arm 1 (n=22), intervention Arm 2 (n=22) or a Control arm (n=22). Following discussion with the farmers, it became apparent that replacement of medicated feed as initially planned for Arm 2 would not be acceptable; therefore, the two intervention arms were merged into one single arm. At the time of the onset of the intervention (May 2018), of 44 farms initially allocated to the intervention, only 31 remained in business; of the 22 allocated to the control, only 8 were still raising chickens. To compensate for the reduced sample size and associated loss in study power, we further allocated four randomly-selected control farms to the intervention arm. Therefore, a total of 35 and 4 farms allocated to the intervention and control arm, respectively, proceeded to the intervention phase (Figure 1).

The intervention commenced with the delivery of the Farmer Training Programme (FTP) in May 2018 to owners of the 35 intervention farms; however, at that time 18 had already restocked with day-olds. Since flocks (n=22) in these farms were not exposed to all four advisory visits, they were therefore analysed as ‘transition’ flocks. Four farms assigned to the intervention arm stopped raising chickens shortly after having attended the FTP modules, and were classified as ‘Baseline-Transition-Stop’ farms.

Data collected from 35 farms (31 intervention, 4 control) were eligible for the final analyses. One hundred flock cycles were analysed as baseline phase (87 in intervention; 13 in control arms) and 89 flock cycles corresponded to the intervention phase (77 intervention farms; 12 in control farms). Of the 77 flocks, 28 (14 farms) were given Product A (an essential oil); and 43 (14 farms) were given Product B (a yeast fraction-based product). Six flocks (3 farms) did not agree with the supplementation of either Product A or Product B.

The median number of chickens restocked per flock was 303 [IQR (inter-quartile range) 200-500], and the median duration of one production cycle was 18 weeks [IQR 16-20]. Each farm raised a median of 5 flocks [IQR 4-7], 2 [IQR 1.0-2.2] during the baseline and 2 [IQR 1.0-2.5] during the intervention phase. Details of number of flocks per farm and status are shown in Table 1 and Figure 2. Descriptive characteristics of chicken farms by total farms, flocks and weeks were presented in Supplementary file 2.

**Table 1.** Categorization of farms based on number of flocks investigated during the baseline phase, transition period and intervention phase.

| Farm group | No. farms (%) | No. flock cycles by status(%) |  |  |  |  |
| --- | --- | --- | --- | --- | --- | --- |
|  |  | Baseline | Transition | Intervention | Control | Total |
| Baseline-Transition-Stop | 4 (3.9%) | 9 | 4 | - | - | 13 (3.9%) |
| Baseline-Transition-Intervention* | 14 (13.7%) | 42 | 18 | 37 | - | 97 (29.4%) |
| Baseline-Intervention* | 17 (16.7%) | 45 | - | 40 | - | 85 (25.8%) |
| Baseline-Control* | 4 (3.9%) | 13 | - | - | 12 | 25 (7.6%) |
| Baseline-Stop | 63 (61.8%) | 110 | - | - | - | 110 (33.3%) |
| Total | 102 (100%) | 219 | 22 | 77 | 12 | 330 (100%) |

\*Data used in further statistical modelling.

**Figure 2.**
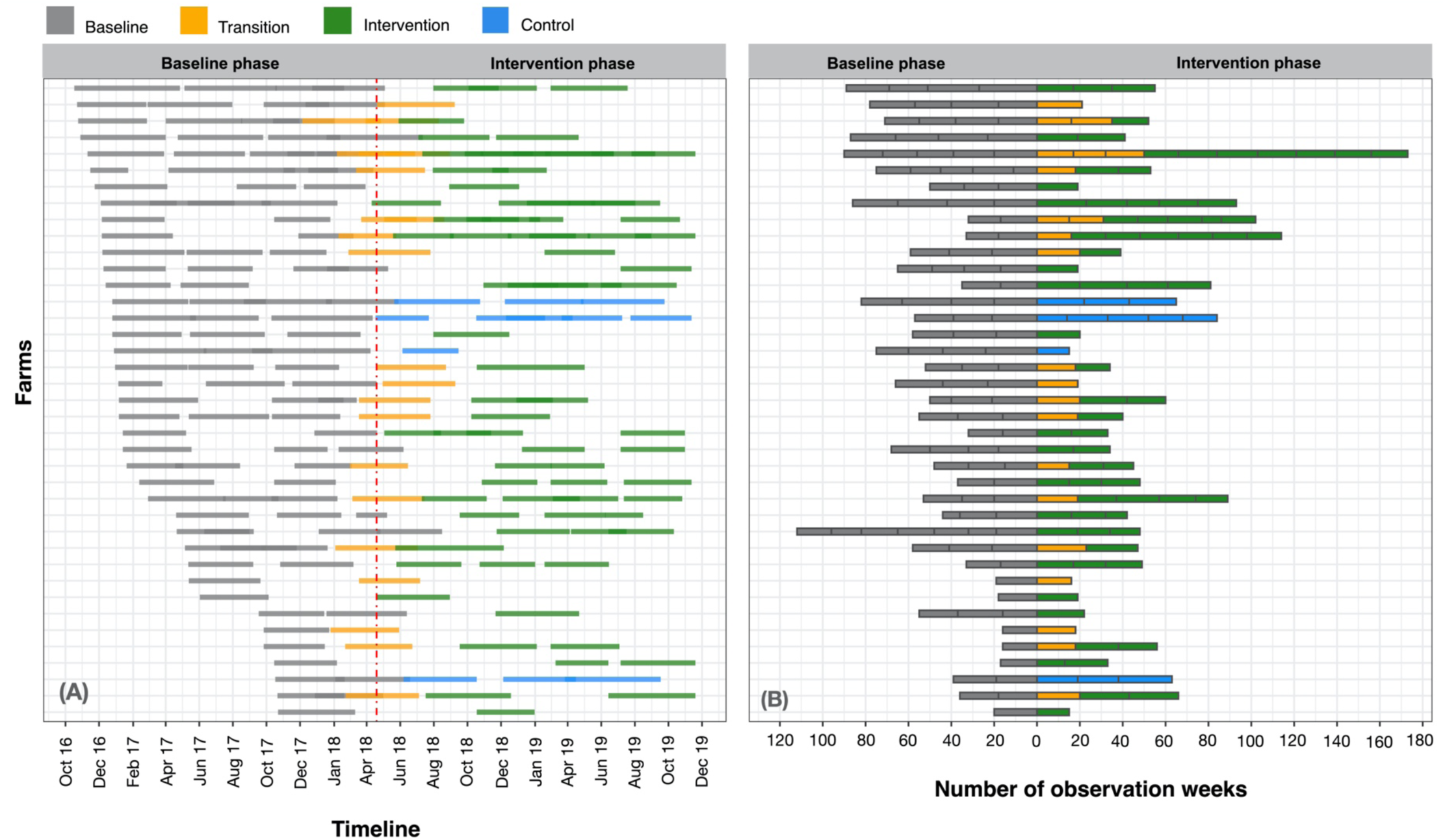
Calendar and observation time of the 39 farms recruited during the baseline and intervention phases. These included: Baseline-Transition-Stop farms (=4), Baseline-Control farm (n=4), Baseline-Transition-Intervention (n=14) and Baseline-Intervention farms (n=17). A total of 63 farms (61.8% of recruited farms) stopped raising chickens and are not displayed in the graphs.

### AMU, mortality and bodyweight of chicken flocks

We collected data over 5,872 weeks; of which 3,899 (66.4%) corresponded to the baseline and 1,973 (33.6%) to the intervention phase. The latter included 396 (6.7%) weeks from transition flocks, 1,350 (23.0%) weeks of full intervention flocks, and 277 (3.9%) weeks from flocks allocated to the control arm. Data on AMU, mortality and bodyweight in these flocks over the baseline, transition and intervention cycles are presented in Table 2.

**Table 2.**
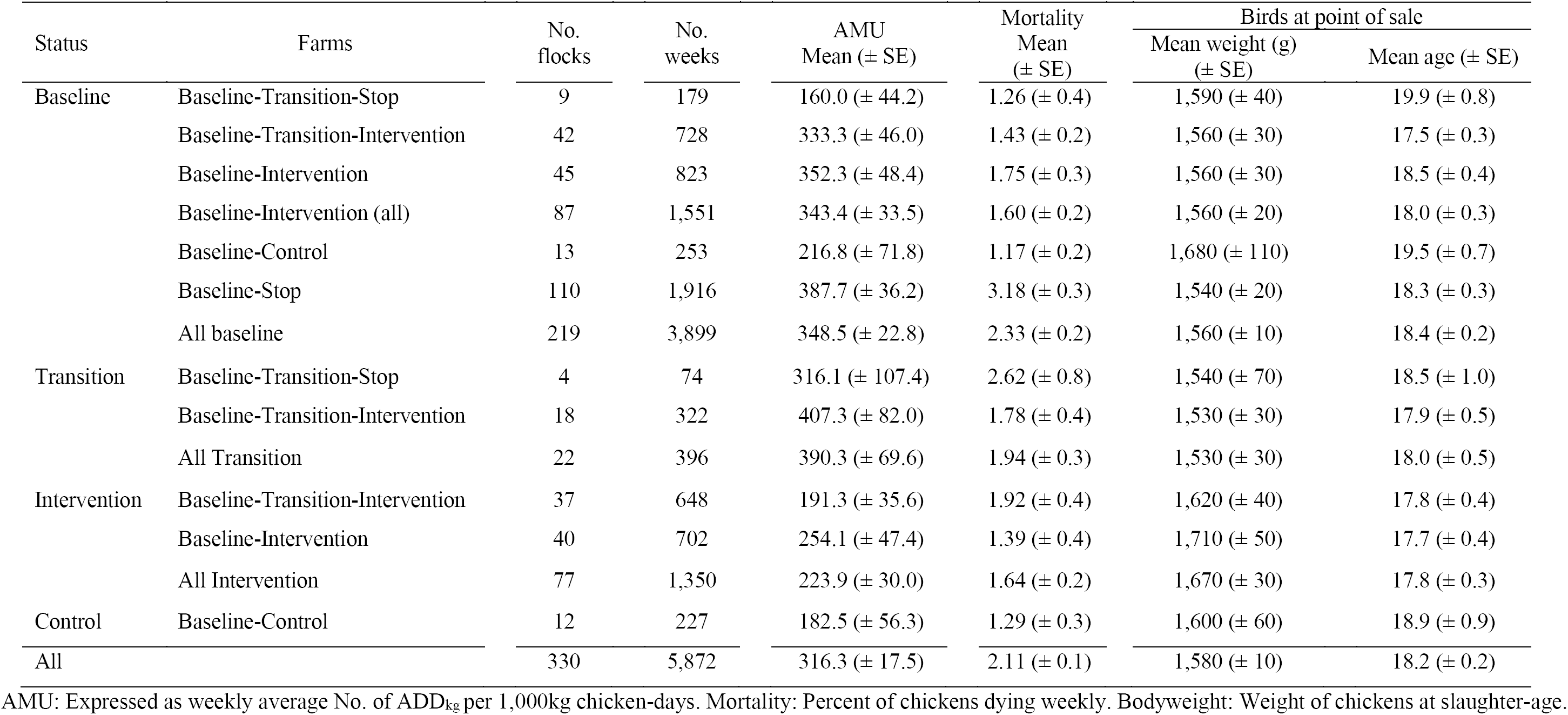
Description of number of AMU, mortality, bodyweights of chicken flocks.

**Table 3.** Mixed regression models investigating the effectiveness of the intervention on AMU, mortality and chicken bodyweight.

| Models | Weekly ADD<sub>kg</sub><sup>†</sup> |  |  | Weekly mortality<sup>††</sup> |  |  | Chicken bodyweight<sup>†††</sup><br>(unit=100g) |  |  |
| --- | --- | --- | --- | --- | --- | --- | --- | --- | --- |
|  | HR | 95% CI | p-value | HR | 95% CI | p-value | β | 95% CI | p-value |
| **Intervention Arm** |  |  |  |  |  |  |  |  |  |
| *Univariable* |  |  |  |  |  |  |  |  |  |
| Thap Muoi district<br>(baseline=Cao Lanh) | 0.68 | 0.37-1.26 | 0.220 | 1.60 | 1.13-2.26 | 0.007 | 0.73 | -0.30-1.77 | 0.179 |
| No. of restocked chickens (log) | 0.55 | 0.36-0.85 | 0.007 | 1.34 | 1.02-1.47 | 0.032 | -0.08 | -0.78-0.65 | 0.820 |
| Status (baseline=Baseline) |  |  |  |  |  |  |  |  |  |
| Transition | 0.77 | 0.30-2.01 | 0.598 | 1.38 | 0.76-2.52 | 0.294 | -0.46 | -1.49-0.58 | 0.390 |
| Intervention | 0.33 | 0.17-0.65 | 0.001 | 0.57 | 0.40-0.82 | 0.002 | 1.00 | 0.37-1.64 | 0.002 |
| *Multivariable 1* |  |  |  |  |  |  |  |  |  |
| Thap Muoi district<br>(baseline=Cao Lanh) |  |  |  | 1.61 | 1.14-2.29 | 0.007 |  |  |  |
| No. of restocked chickens (log) | 0.55 | 0.36-0.85 | 0.007 | 1.39 | 1.07-1.81 | 0.015 | -0.05 | -0.72-0.65 | 0.884 |
| Phase (baseline=Baseline) |  |  |  |  |  |  |  |  |  |
| Transition | 0.86 | 0.34-2.18 | 0.743 | 1.35 | 0.74-2.45 | 0.327 | -0.45 | -1.49-0.59 | 0.393 |
| Intervention | 0.34 | 0.18-0.66 | 0.002 | 0.60 | 0.41-0.86 | 0.005 | 1.00 | 0.37-1.64 | 0.002 |
| *Multivariable 2* |  |  |  |  |  |  |  |  |  |
| Thap Muoi district<br>(baseline = Cao Lanh) |  |  |  | 1.62 | 1.14-2.30 | 0.007 |  |  |  |
| No. of restocked chickens (log) | 0.55 | 0.36-0.85 | 0.007 | 1.39 | 1.07-1.81 | 0.015 | -0.06 | -0.73-0.63 | 0.842 |
| Phase (baseline=Baseline) |  |  |  |  |  |  |  |  |  |
| Transition | 0.86 | 0.34-2.18 | 0.743 | 1.35 | 0.74-2.45 | 0.327 | -0.46 | -1.48-0.60 | 0.404 |
| First intervention cycle | 0.46 | 0.20-1.05 | 0.066 | 0.57 | 0.36-0.92 | 0.022 | 0.77 | -0.04-1.59 | 0.068 |
| Subsequent intervention cycle | 0.26 | 0.10-0.63 | 0.003 | 0.61 | 0.40-0.94 | 0.025 | 1.19 | 0.44-1.97 | 0.003 |
| **Control Arm** |  |  |  |  |  |  |  |  |  |
| *Univariable* |  |  |  |  |  |  |  |  |  |
| Thap Muoi district<br>(baseline = Cao Lanh) | 0.95 | 0.15-6.13 | 0.958 | 1.88 | 0.49-7.32 | 0.360 | 1.49 | -3.74-6.71 | 0.672 |
| No. of restocked chickens (log) | 0.19 | 0.04-0.87 | 0.033 | 0.08 | 0.04-0.17 | <0.001 | 1.26 | -2.01-4.49 | 0.471 |
| Status (baseline=Baseline) |  |  |  |  |  |  |  |  |  |
| Intervention calendar time | 1.07 | 0.18-6.23 | 0.937 | 0.60 | 0.17-2.09 | 0.419 | -0.02 | -2.72-2.23 | 0.983 |
| *Multivariable 3* |  |  |  |  |  |  |  |  |  |
| Thap Muoi district<br>(baseline=Cao Lanh) | 0.64 | 0.09-1.73 | 0.666 | 0.53 | 0.22-1.27 | 0.156 | 2.33 | -3.19-7.88 | 0.594 |
| No. of restocked chickens (log) | 0.15 | 0.03-0.87 | 0.034 | 0.07 | 0.04-0.15 | <0.001 | 1.73 | -1.65-4.65 | 0.383 |
| Phase (baseline=Baseline) |  |  |  |  |  |  |  |  |  |
| Intervention calendar time | 1.42 | 0.24-8.44 | 0.696 | 0.81 | 0.38-1.74 | 0.591 | 0.14 | -2.74-2.21 | 0.905 |

Multivariable 1 intercepts: <sup>†</sup>-11.887 (SE=1.303); <sup>††</sup>-11.400 (SE=0.794); <sup>†††</sup>16.093 (SE=1.908). Multivariable 2 intercepts: <sup>†</sup>-11.897 (SE=1.308); <sup>††</sup>-11.400 (SE=0.793); <sup>†††</sup> 16.093 (SE=1.908). Multivariable 3 intercepts: <sup>†</sup> -5.135 (SE=5.539); <sup>††</sup> 7.382 (SE=2.297); <sup>†††</sup> 4.558 (SE=12.129).

During the baseline phase, flocks (n=110) raised in the 63 farms that dropped out prior to the implementation of the intervention phase had a higher mortality (weekly average 3.18 per 100 birds; SE±0.3), than flocks (n=109) in 39 farms that proceeded to the intervention (1.52 per 100 birds; SE±0.1) (Wilcoxon Test, p=0.020).

The weekly summary data of the outcome variables and the distribution of flocks concerning these in flocks during the baseline (n=87) and intervention phases (n=77) are displayed in Figure 3. Weekly AMU in these flocks was reduced from 343.4 (SE±33.5) (baseline) to 223.9 (SE±30.0) (intervention) Animal Daily Doses (ADD_kg_) per 1,000 kg chicken-days (−34.8%) (one-sided Wilcoxon test, p<0.001). The bodyweight at slaughter-age of chickens of intervention flocks was 1,670 g (SE±30), compared with 1,560 g (SE±20) during baseline (+7.1%) (one-sided Wilcoxon test, p=0.006). However, weekly mortality increased from 1.60 (per 100 birds) (SE±0.2) to 1.64 (SE±0.2) (+2.4%), although the difference was not significant (one-sided Wilcoxon test, p=0.999).

**Figure 3.**
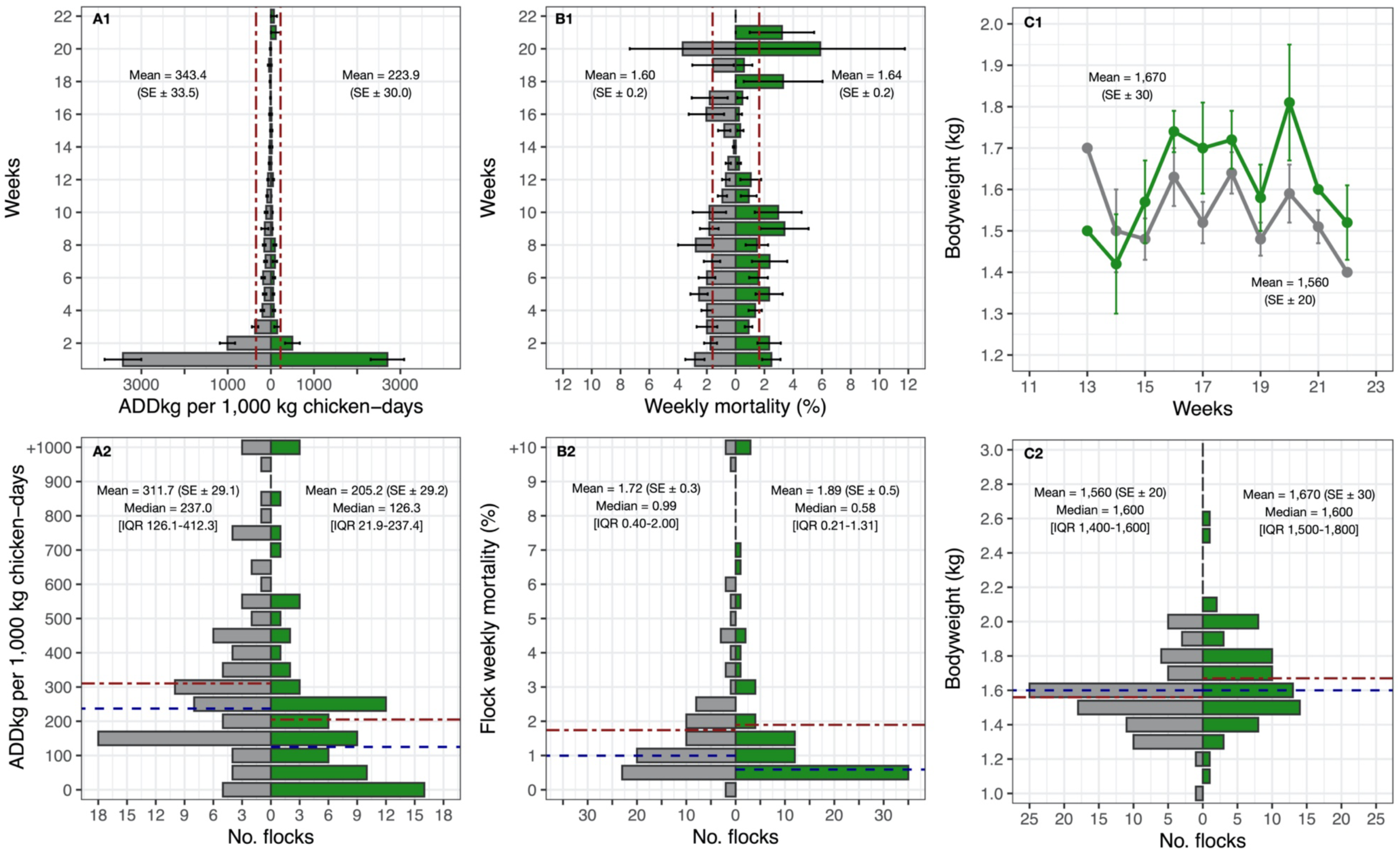
Comparisons between AMU, mortality and bodyweight by weeks and flocks between baseline (in gray colour) and intervention phase (in green colour) in 31 eligible farms. Dot-dash red line: Mean. Dashed blue line: Median.

The unadjusted overall mortality increased slightly during the intervention. However, the number of farms that experienced a reduction in mortality exceeded (19/31) than those that increasing it (12/31). The changes in (flock average) values of ADDkg per 1,000 kg chicken-days, mortality and bodyweight between the baseline and intervention phases are displayed in Figure 4. Among intervention flocks, there were 3/77 (3.9%) with an average weekly mortality greater than 12% (12.8%, 24.8% and to 26.0%) and a cumulative mortality of >98%; two of these flocks were detected with Highly Pathogenic Avian Influenza (HPAI) and one with *Avibacterium paragallinarum*, compared with 2/87 flocks experiencing >10% weekly mortality among baseline flocks (one 12.4% and one 12.8%) and cumulative mortality of 100% in these two baseline flocks.

**Figure 4.**
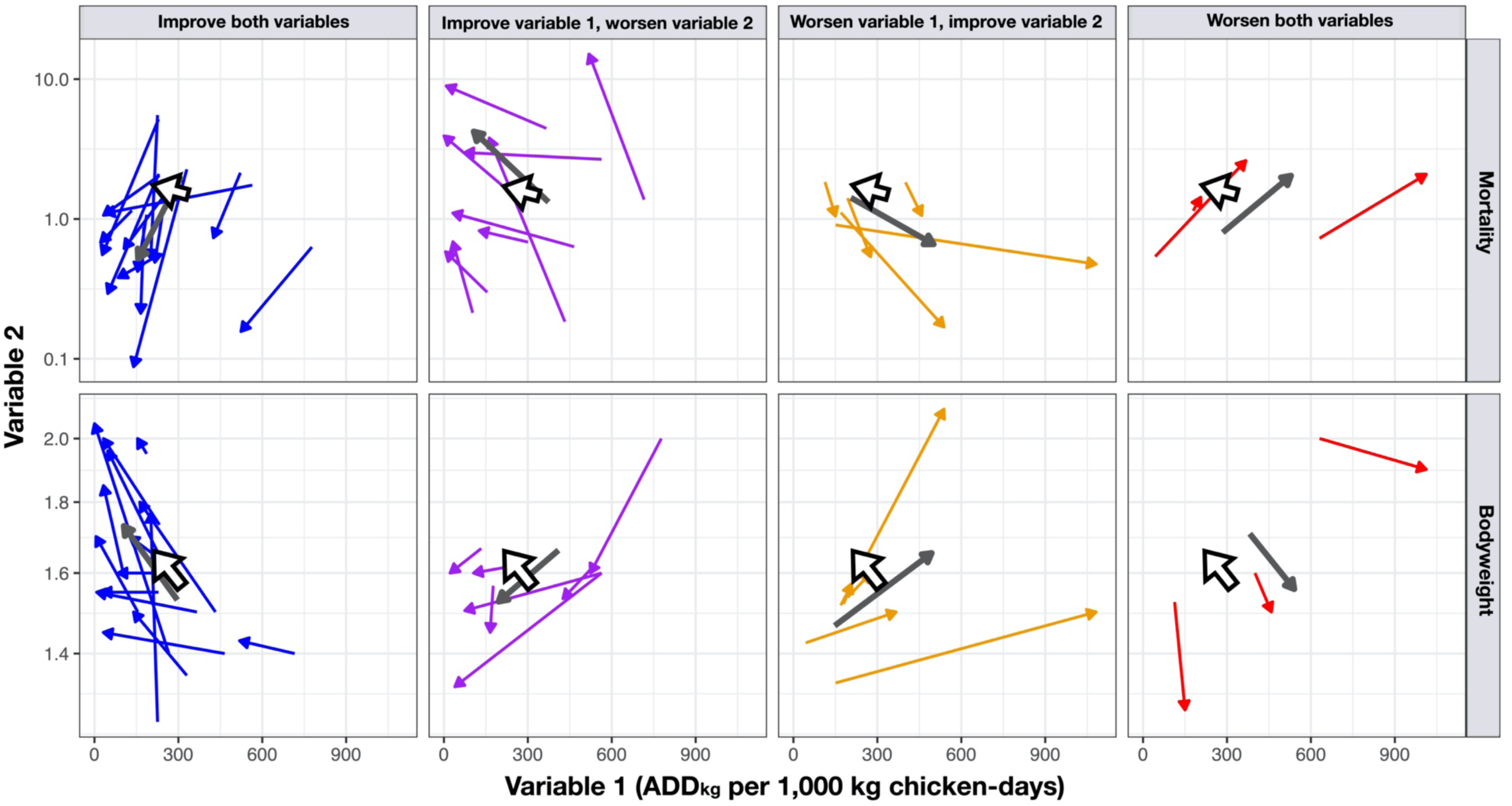
Bi-dimensional arrow charts showing crude (unadjusted) changes between the baseline and the intervention phases with regards to Variable 1 (No. ADDkg per 1,000 kg chicken-days) and Variable 2 (mortality and bodyweight). The arrow indicates the direction of change. Large gray arrows indicate summary result for each pair of variables. Large white arrows indicate overall results. The log10-transformed scale for easier visualization.

In the four farms that were allocated to the control arm, a total of 13 flocks were investigated during the baseline phase, and 12 during the intervention phase. AMU in these decreased from 216.8 (SE±71.8) to 182.5 (SE±56.3) (Wilcoxon Test, p=0.857); weekly mortality changed from 1.17 to 1.29 per 100 birds (Wilcoxon test, p=0.493) and bodyweight changed from 1,680 g to 1,600 g (Wilcoxon test, p=0.511).

### Correlation between AMU and mortality, bodyweight

There were significant correlations between weekly AMU (ADD_kg_ per 1,000 kg chicken-days) and mortality (Spearman’s rank correlation R=0.26; p<0.001). There were, however, no correlations between average bodyweight and AMU (R=-0.06, p=0.292) or mortality (R=-0.09, p=0.128) at flock level. The details of these calculations are provided in Supplementary file 3.

### Modelling

In the univariable models for the Intervention Arm, ‘Status=intervention’ was associated with an overall decreased AMU (HR=0.33; 95% CI=0.17-0.65; p=0.001) (−67%), decreased mortality (HR=0.57; 95% CI=0.40-0.82; p=0.002) (−43%) and increased bodyweight (+100g; 95% CI 37-164 p=0.002). The size of the flock was negatively associated with AMU (HR=0.55, 95% CI 0.36-0.85, p=0.007), but positively associated with mortality (HR=1.34; 95% CI 1.02-1.47; p=0.032). Adjustment for flock size resulted in minimal change in the estimates of AMU (−66%) (HR=0.34; 95%CI 0.18-0.66; p=0.002), mortality (−40%) (HR=0.60; 95% CI 0.41-0.86; p=0.005) and bodyweight (+100g; 95% CI 37-164g; p=0.002). When the variable level ‘status=intervention’ was replaced by two new variables (first, subsequent cycles), greater reductions in AMU were seen in the subsequent (HR=0.26, p=0.003) compared with the first cycle (HR=0.46; p=0.066), although the difference between both was not statistically significant (p=0.298). Similarly, chicken bodyweight further increased during subsequent intervention cycles (+119g per chicken sold, p=0.003) compared with the first intervention one (+77g, p=0.068) (p=0.378). Levels of mortality did not change between first and subsequent cycles (p=0.967). There were no significant interactions between either ‘flock size’ and ‘district’ and ‘Status=intervention’. There was no statistical difference between in AMU and mortality between flocks using Product A, Product B or those given no additional product. However, flocks that were administered with either Product A and B had increased bodyweight compared with flocks not given any supplementary product (data not shown).

In the control arm, there were no significantly associations between ‘Status=intervention calendar time’ and any of the three outcome variables in either univariable or multivariable models (all p>0.419). After adjustment of flock size and study district in multivariable models, estimates of AMU and bodyweight increased (+42% and +14g, respectively) and mortality was reduced (−19%).

## Discussion

Through a locally delivered veterinary intervention, we achieved a 66% reduction in antimicrobials (quantified as daily doses) administered to small-scale commercial chicken flocks, alongside a reduction in mortality (−40%). In our crude (unadjusted) analyses AMU reductions were, however, modest (−35%), since our analysis implicitly adjust for week of use and most AMU took place during the early weeks (i.e. the brooding period). Similarly, the crude data indicated a slightly higher mortality during the intervention (+2.4%). However, the adjusted analysis indicated a ∼40% reduction in mortality, and mortality was reduced in a majority (19/31) of farms. This discrepancy was explained unusually high mortality in three intervention flocks.

Unlike other studies involving the delivery of a uniform treatment (i.e. vaccination) (Bessell et al., 2017), our intervention consisted of providing farmers with veterinary advice. The nature of this advice was variable across farms, and was based on specific observations and information collected by project veterinarians from their flocks. This advice included measures to improve flock health and productivity, whilst emphasizing the message that ‘antimicrobials should not be admnistered to healthy chickens’.

In addition to providing antimicrobial replacement products, the main advice given to farmers focused on biosecurity, cleaning and disinfection, vaccination, litter management and administration of medicines (including antimicrobials, antiparasitic drugs and other health-enhancing products). The detail of this advice provided and its uptake will be presented elsewhere. The advice provided was based on a persuasive, rather than a restrictive advice. We believe this approach is likely to be more sustainable in the mid-to-long term (Davey et al., 2013). A similar holistic approach was adopted on a study on pig farms in Belgium, resulting in 52% AMU reduction in pigs raised from birth to slaughter, and by 32% among breeding animals; furthermore, the study resulted in additional productivity gains (Postma et al., 2017). Similarly, a study conducted in four European Union (EU) countries reported AMU reductions of 3% and 54% in fattening in weaned pigs, respectively following improvements of herd management practices (Raasch et al., 2020). However, reductions in AMU were not seen in breeding pigs, and the authors attributed it to the concurrent incursion of Porcine Epidemic Diarrhoea (PED) in Germany. A study in 20 industrial-scale broiler farms in Europe using a holistic approach resulted in 20% reductions in levels of AMU and 14% increase in gross margins (Roskam et al., 2019).

After consultation with participating farmers during the baseline phase, we were compelled to modify our original protocol by offering selected health-enhancing, antimicrobial replacement products (Talkington et al., 2017) to chicks during the brooding phase. This aimed at allaying the farmers’ anxiety about reducing or eliminating antimicrobials during this critical phase of production. Administration of antimicrobials during the brooding phase is standard practice and many antimicrobial-containing commercial formulations are marketed as ‘brooding medicine’ (Carrique-Mas et al., 2019). Similarly, many of our study farmers expressed their opposition about changing the feed and therefore we consolidated the two intervention arms into only one arm. Often the advice provided by project veterinarians to farmers was overrun by that given at local vet drug shops. Farmers often visit these shops to buy animal feed and other supplies (Phu et al., 2019). In addition, the antimicrobial product labels often include indications for prophylactic use at a lower dose (Yen et al., 2019).

Small-scale commercial chicken production using native breeds is widespread in the Mekong Delta of Vietnam, and often represents an upgrade from backyard production. The popularity of this system resides in the preference of the Vietnamese consumer for meat of long-cycle native birds. Native chicken meat reaches a considerably higher price compared with broiler meat (PW, 2018). However native chickens (and their crosses) are slow growing (>4 months), and preventing disease over such a prolonged period requires sustained efforts (Carrique-Mas et al., 2019).

In our study, the identification and enrollment of study farms was challenging due to the fluidity of this type of production system, with many households setting up chicken farms as well as stopping raising chickens altogether. Because of this, a large number of farms did not remain in business over the extended duration of the study. Indeed, flock mortality was an important predictor for farmers giving up raising chickens (data not shown) and a large fraction of our study farms (61.8%) had gone out of business even before the start of the planned intervention phase.

In addition to their previous experience with disease, farmers may start or stop raising chickens depending on circumstances, such as market price of day-olds, commercial feed and poultry meat, income from the sale of the previous flocks or other rural activities. Furthermore, many farmers raised one cycle per year, but not necessarily every year. This was reflected in the lack of experience in chicken husbandry of many farmers (and farm workers). This represents a hurdle for the implementation of correct management practices. This contrasts with a recent study in Belgium, where pig farmers had on average 22.6 years of experience (Postma et al, 2016). In this context, often antimicrobials are used as replacement of other, most costly, but demanding husbandry practices (Truong et al., 2019). The incursion of African Swine Fever (ASF) in Vietnam in January 2019 and its spread within the country (VASFU, 2019) coincided with the intervention phase in this study. This may have exerted additional pressures over our study farmers. During this time, many farms in ASF-affected provinces switched to chicken production, resulting in increased market availability of low-cost chicken meat, therefore reducing the value of chicken production in our area.

The changes to the initial study design are a testament to the challenges of conducting intervention studies in small-scale farming systems. Initially, we planned to allocate one third of all recruited farms to the control arm in order to measure any environmental influences on AMU, for example, due to public engagement initiatives (television campaigns, work in schools, etc.) that took place in the province under the umbrella of this project. Exposure to these may have inadvertently had an influence on the farmers’ decision on AMU beyond the intervention. Given the high number of farms that stopped chicken production, we opted for reducing the size of the control arm to a minimum of four, thus reducing the statistical power of any analysis in that group. However, the descriptive data from this small control group suggests no change between baseline and intervention, and gives additional validity of the observed findings.

The study demonstrates that reducing current high levels of AMU through the provision of veterinary advice is achievable in the Vietnamese small-scale commercial farming context. There was an indication that farmers responded to the advice given, and supplementation with health-enhancing products may be beneficial. Many farmers, especially the larger ones may even be willing to pay for such a service, since labour costs in Vietnam are relatively low (approximately 25 USD for a two-hour visit). We propose to develop a business case for an advisory service targeting the main livestock-producing regions in the country (Mekong River Delta, Southeast, Central region, Red River Delta), with the value proposition that healthy livestock means profitable businesses.

## Materials and Methods

### Study design

The intervention was designed as a randomized ‘before-and-after’ controlled study on farms raising chickens for meat in two districts (Cao Lanh and Thap Muoi) within Dong Thap province (Mekong Delta, Vietnam) (Figure 1 and Supplementary file 1). The study was designed in two stages, a ‘baseline’ followed by an ‘intervention’ phase. Two intervention arms (Arm 1 and Arm 2) were initially planned, both including the provision of training and advice to farmers, as well as a control arm (Arm 3) (no training or advice). The difference between both intervention arms was that Arm 2 also included the withdrawal of medicated commercial feed. This aimed at investigating whether restriction of medicated feed might have affected disease outcomes, therefore contributing to changes in levels of AMU (Carrique-Mas & Rushton, 2017).

Farmers registered in the official SDAH census (2014) were contacted by post and were invited to participate in an introductory meeting held in October 2016 in each of the two study districts. In these meetings, the project aims and methods were outlined. Farmers willing to enroll in the study were asked to contact project staff as soon as they restocked with day-old chicks. Farmers that restocked with chicken flocks (defined as a group of birds raised together in the same building) meeting the criterion ‘>100 meat chickens raised as single age’ were enrolled.

### Description of the baseline and the intervention

During the baseline phase of the study, routine AMU and productivity data were collected from enrolled farms without the provision of any advice. Using a random number generator, we allocated enrolled farms to either an intervention or a control arm. All farms allocated to the intervention arm were supported with a Farmer Training Programme (FTP), where farm owners were invited to participate in six workshops where a poultry veterinarian instructed them on the principles of chicken husbandry, prevention, control of infectious diseases and waste management and a Farm Health Plan (FHP), where each farm was assigned to a Project Veterinarian (PV) who was responsible for providing specific advice to farmers. The PV visited each farm on three different occasions for each flock cycle: (i) early-brooding (weeks 1-2), (ii) late brooding (weeks 3-4), and (iii) grow-out (>2 months) periods. Prior to each visit, the PV reviewed records of productivity and disease over previous cycles, inspected the flock and house/pen, reviewed farmers’ records, discussed with farm owner about current production/health issues, and then drafted a list of recommendations to address them. In addition, the PV proposed the farm owner the use of an antimicrobial replacement product, either an essential oil-based (Product A) or a yeast fraction-based product (Product B) used 3 days/week over the first 10 weeks of the production cycle. The allocation of either of these products was based an assessment of the history of disease in the flock (Product A) if the farm had a history of diarrhoea in previous flocks; or Product B, all other flocks. In all visits, the PVs reminded the farmers that healthy birds should not be given any antimicrobials.

### Data collection

Each farmer was provided by project staff with a diary to weekly record data on farming practices, including number of chickens purchased, number of chickens in and out of the flock (number of dead and sold chickens), as well as the types and quantities of antimicrobial products used. The average bodyweight of slaughter-age chickens was also measured by average of total bodyweight of chickens divided for total number of chickens sold. Project staff visited study farms four times (different from PV visits) to verify the data collected, which was subsequently transferred to validated questionnaires and double-entered into a web-based database.

### Statistical analyses

The initially proposed sample size was based on previous quantitative data on AMU in Mekong Delta chicken farms (Carrique-Mas et al., 2015). We aimed at recruiting 120 farms and estimated a total of 40 farms for each arm. A sample size of 40 farms per arm, each contributing with 2 cycles investigated during baseline and 2 during the intervention, and a two-sided significance level of 5%, will have 82% power to detect a ∼33% reduction, and a 91% power to detect a 50% reduction. Since the study design exploits within-farm correlation of unknown magnitude, the true power was expected to be higher.

The primary outcome was the weekly number doses of antimicrobial active ingredient (AAI) corresponding to 1 kg of live chicken administered to a flock (Weekly ADD_kg_). Secondary outcomes were ‘Weekly mortality’, calculated by dividing the number of chickens dying over the week by the total chicken present at the beginning of each week (%), and ‘Weight of the birds (in units of 100g) at the time of sale’. The latter was calculated by dividing the total flock weight by the number of chickens sold at the end of the cycle. The correlation between all three outcomes at flock and at week level was investigated using the Spearman’s rank correlation coefficient.

Weekly antimicrobial consumption (ADD_kg_) was calculated by multiplying the amounts of antimicrobial product administered by the farmer to the flock (g) through water or feed, multiplied by a dilution factor (DF) indicating the ratio of product to water/feed. The DF indicated the amount of product to be diluted in water (Volume of water /Weight of product) (l/g) or mixed with feed (Weight of feed /Weight of product) (kg/g) based on the product label. The obtained amounts were then divided by the estimated daily consumption of water (0.225 l) or feed (0.063 kg) by a 1 kg-chicken (Cuong et al., 2019).

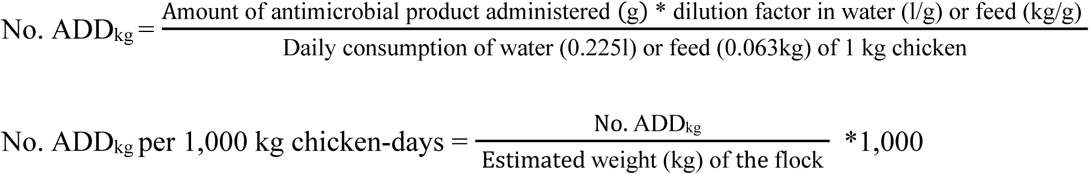

We built Poisson regression models with for ‘Weekly ADD_kg_’ and ‘Weekly mortality’. For the former the offset was the (weekly) total number of chicken-kg days (log); for the latter it was the number of chickens at the beginning of the week (log). In addition, a linear regression model was developed with bodyweight of chickens at the point of sale (kg) as outcome. In all cases, ‘Farm’, ‘Flock cycle’ and “Week” were modelled as random effects, where ‘Week’ was nested within ‘Flock cycle’, and the latter was nested within ‘Farm’. The main variable of interest was the impact of the intervention delivered; therefore, we investigated ‘Status’ (baseline, transition, and intervention) as an explanatory variable in Intervention Arm and ‘Status’ (baseline, intervention calendar time) as an explanatory variable in the Control Arm. ‘Status=transition’ was assigned to those flocks that were not exposed to all three advisory visits for Intervention Arm farms. This occurred to a number of flocks at the beginning of the intervention phase, given that some advisory visits (typically the first and second) were missed. In order to account for the potential confounding effects of ‘District’ and ‘Flock size’ these were forced into a multivariable model; we tested the interactions between ‘Status=intervention’ with ‘District’ and ‘Flock size’ to investigate whether the observed effects were dependent on the geographical location or the size of the flock. Moreover, we investigated whereas subsequent cycles over the intervention resulted in improved outcomes by splitting ‘Status=intervention’ into ‘Status=first intervention cycle’ and ‘Status=subsequent intervention cycle’. The presence of overly influential observations was investigated by testing the model with and without those observations yielding the largest residuals. We used the ‘survey’ package to calculate (farm-flock-week) adjusted estimates and ‘lme4’ package to build statistical models (http://www.r-project.org).

## Acknowledgements

The authors would like to thank all farmer participants, the staff affiliated to the Dong Thap Sub-Department of Animal Health, Production and Aquaculture, the staff at Agricultural Service Center of Cao Lanh and Thap Muoi district, Dong Thap province for their help and support.

## Author contribution

JCM, JR, NVC conceived and designed study; DHP, NVC, DBT conducted field survey; BTK, VBH designed and aided data collection; HTVT, LKY, NTTM, ES aided intervention packages; DHP, JCM, NVC, DBT contributed to data analyses; DHP, NVC, JCM, PP, JR, GT contributed to writing up and editing the manuscript. All authors read and approved the final manuscript.

## Ethics statement

The study was granted ethics approval by Oxford Tropical Research Ethic Committee (OXTREC) for Minimal Risk (Ref.5121/16) and by the People’s Committee of Dong Thap province.

## Additional files

**Supplementary file 1.**
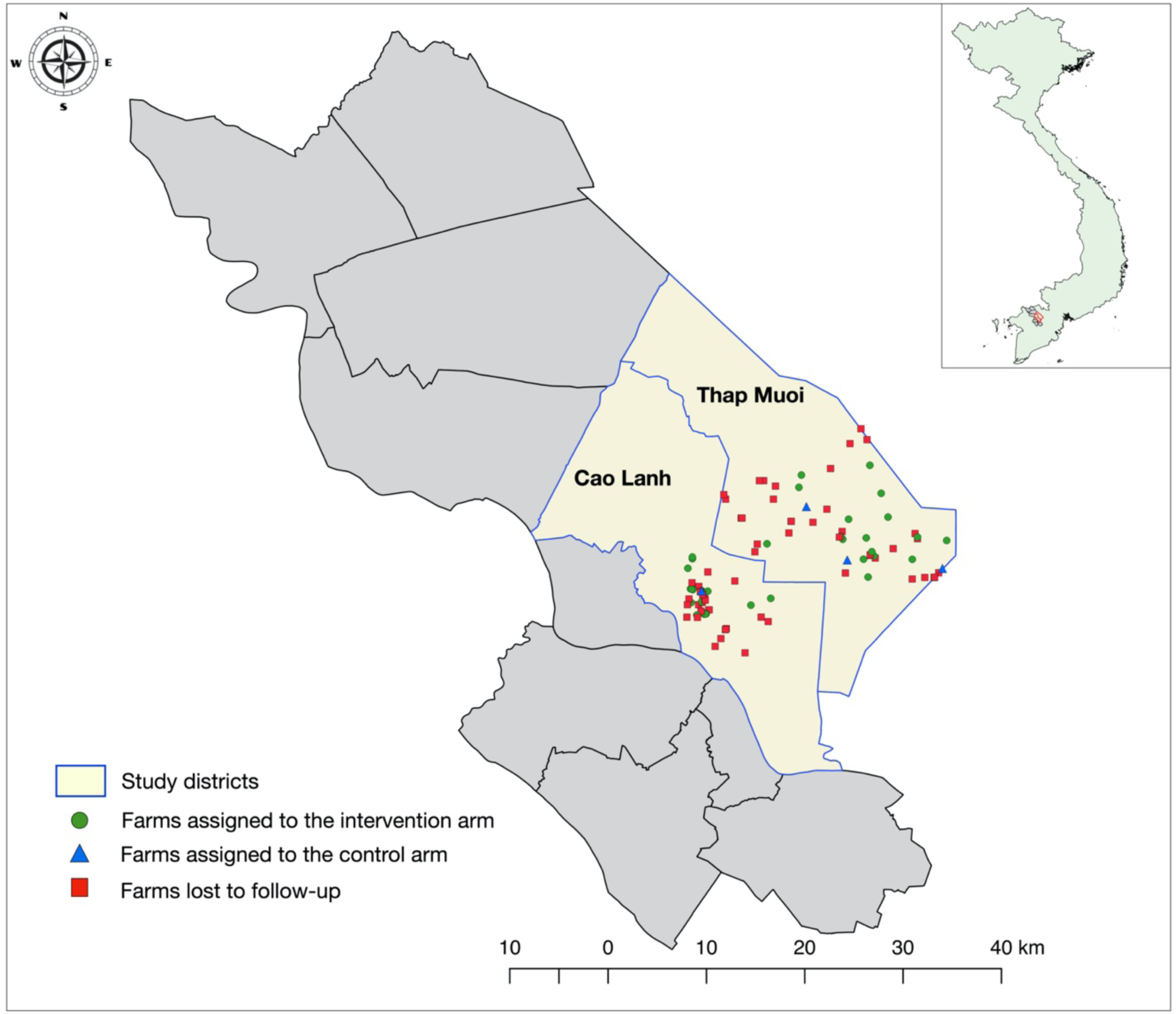
Study area in the province of Dong Thap, Mekong Delta of Vietnam

**Supplementary file 2.**
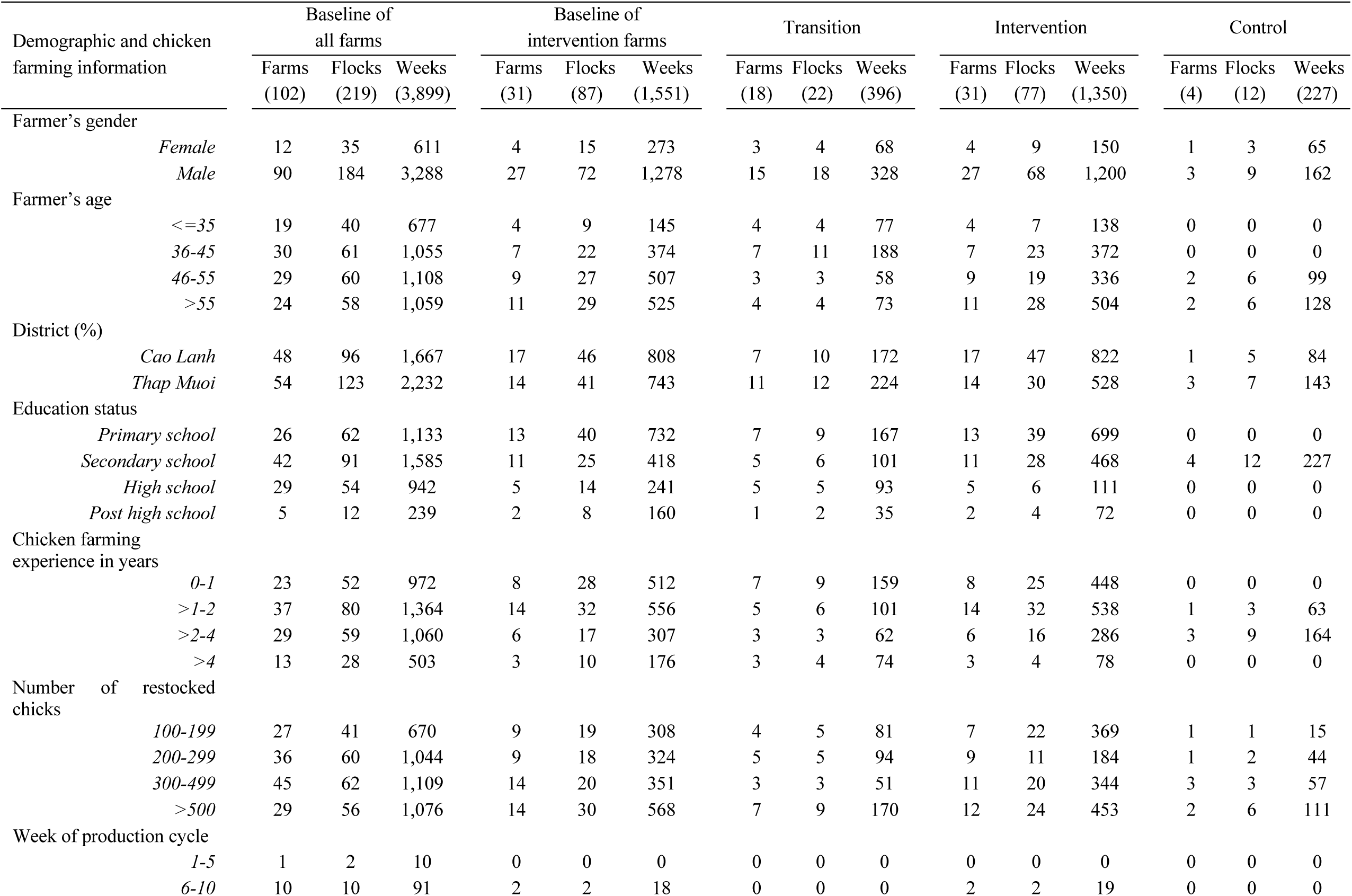
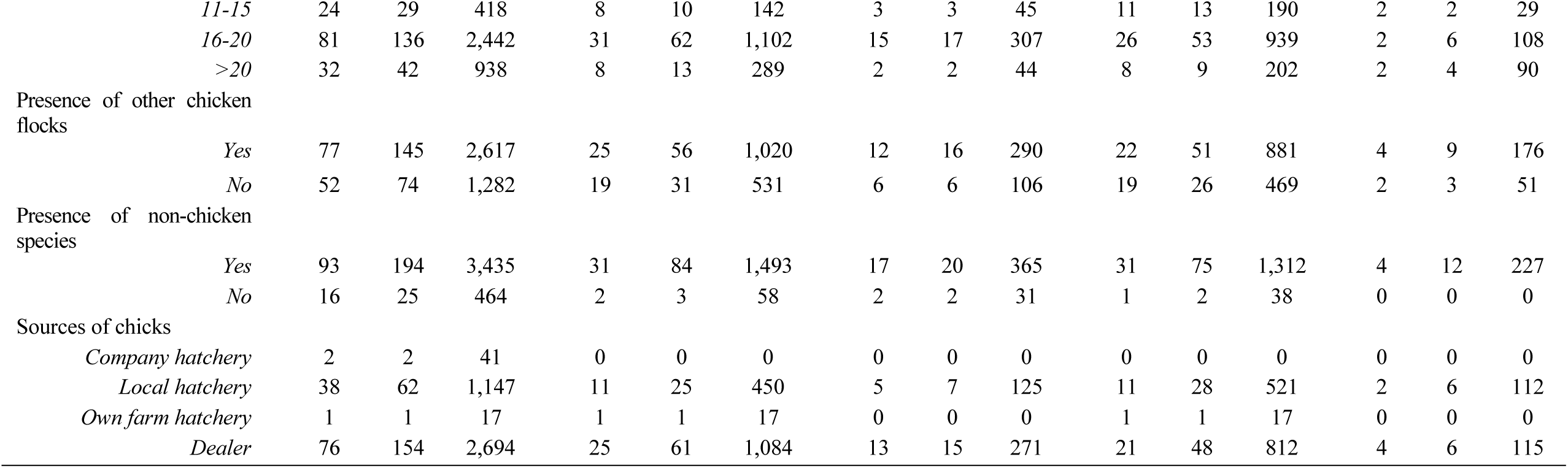
Descriptive characteristics of chicken farms by total of farms, flocks and weeks.

**Supplementary file 3.**
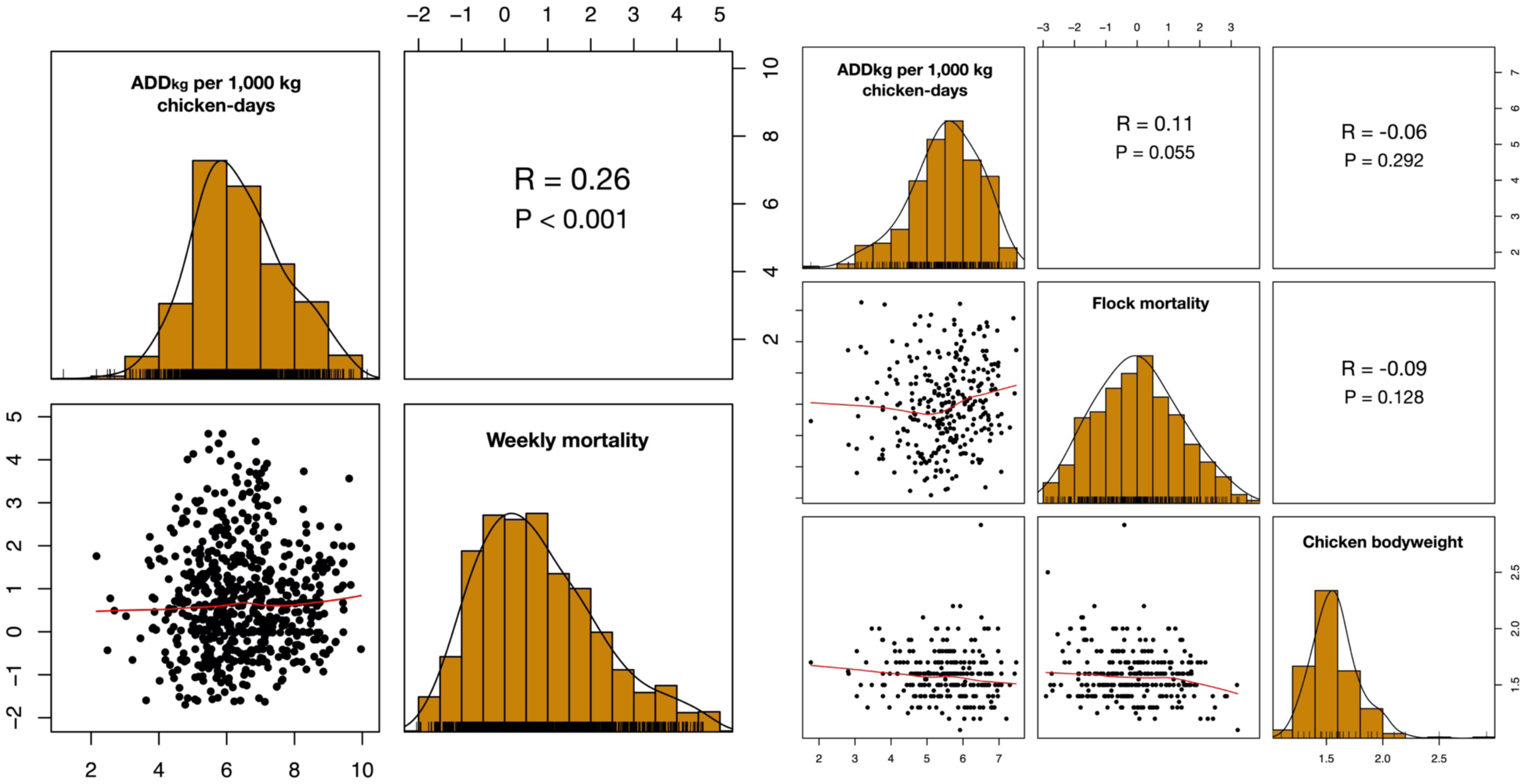
Correlation between AMU, mortality and bodyweight by week (left block) and by flock (right block). The data have been log-transformed scale for easier visualization.

